# Chai-1: Decoding the molecular interactions of life

**DOI:** 10.1101/2024.10.10.615955

**Authors:** Chai Discovery, Jacques Boitreaud, Jack Dent, Matthew McPartlon, Joshua Meier, Vinicius Reis, Alex Rogozhnikov, Kevin Wu

## Abstract

We introduce Chai-1, a multi-modal foundation model for molecular structure prediction that performs at the state-of-the-art across a variety of tasks relevant to drug discovery. Chai-1 can optionally be prompted with experimental restraints (e.g. derived from wet-lab data) which boosts performance by double-digit percentage points. Chai-1 can also be run in single-sequence mode without MSAs while preserving most of its performance. We release Chai-1 model weights and inference code as a Python package for non-commercial use and via a web interface where it can be used for free including for commercial drug discovery purposes.

## 1 Introduction

Understanding the three-dimensional structure of biological molecules is critical for studying how they function and interact [1, 2, 3]. This understanding, in turn, is foundational to designing therapeutic molecules targeting the cellular machinery of life [4, 5]. Over the last few years, significant progress has been made using deep learning methods to predict the folded structures for proteins [6, 7, 8] and nucleic acids [9]. More recently, methods like RoseTTAFold All-Atom [10] and AlphaFold3 [11] have introduced models that can predict a wide range of protein and nucleic acid structures, covalent modifications thereof, and small molecule ligand interactions with these complexes.

Here we introduce Chai-1, a state-of-the-art and openly accessible foundation model for biomolecular structure prediction. We demonstrate that Chai-1 excels on a variety of tasks including protein-ligand structure prediction and protein multimer prediction. Furthermore, while Chai-1 is designed to predict biopolymer structures directly from raw sequence and chemical inputs, it can optionally be prompted with experimental constraints such as those provided by epitope mapping or cross-linking mass spectrometry experiments to achieve even more accurate predictions of difficult binding complexes. Chai-1 performs best when given multiple sequence alignments (MSAs), but can also produce strong predictions without MSAs in single sequence mode. In single sequence mode, Chai-1 outperforms ESMFold, and can even outperform AF-Multimer2.3 [12] – which we evaluate with MSAs – under certain evaluations.

Chai-1 model weights and inference code are available for non-commercial use^1^. We also provide a web server^2^ for interfacing with the model, which is made available for commercial use (including for drug discovery tasks).

## 2 Results

### 2.1 Model Architecture

Our model architecture and training strategy largely follows that of Abramson et al. [11] with the key difference that we train a single model with a training data date cutoff of 2021-01-12 as opposed to training separate models for separate evaluations. We also make a number of key additions to enable new functionality (Figure 1), summarized below.

**Figure 1.**
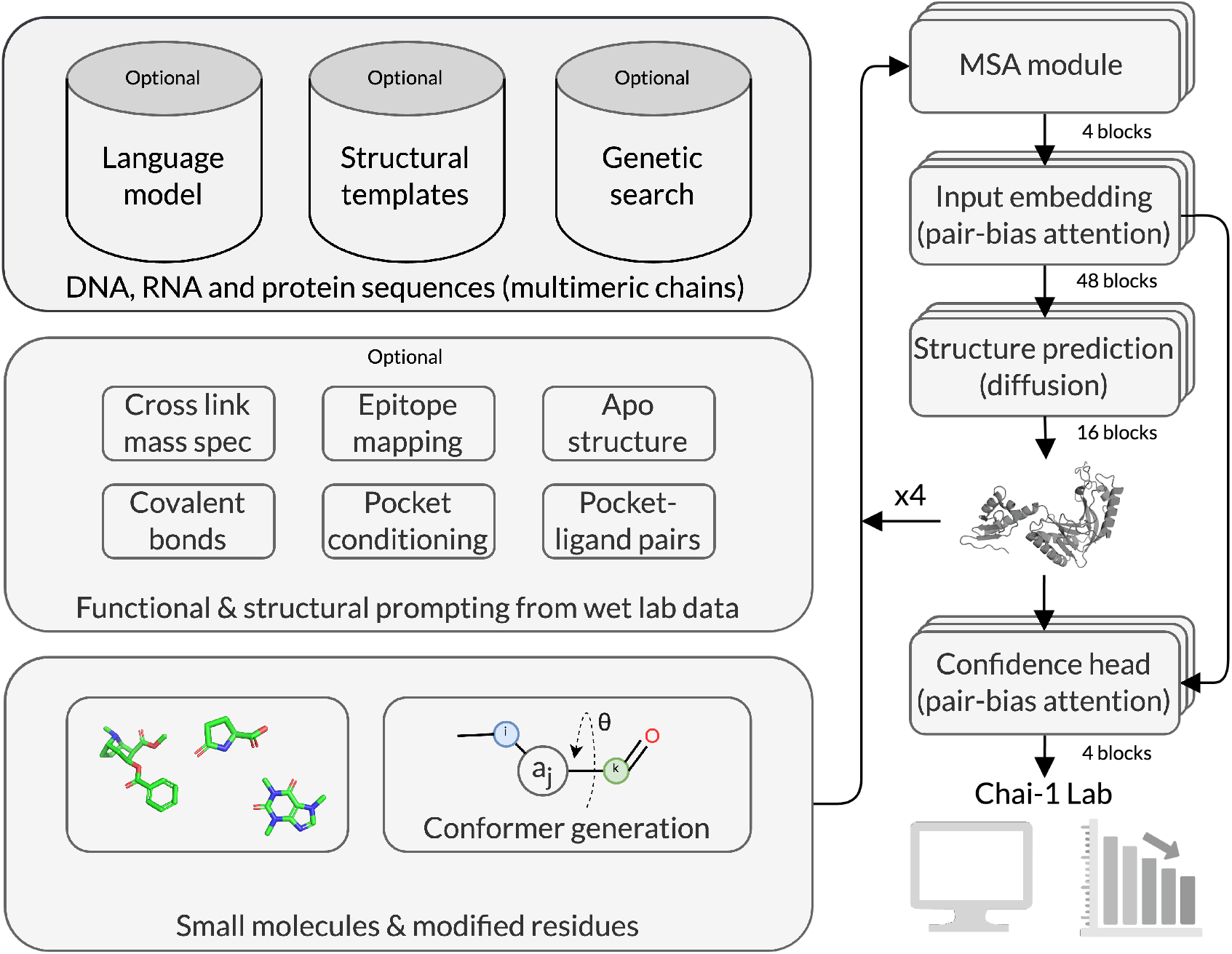
Overview of the Chai-1 model architecture and input features. Chai-1 accepts a wide variety of optional input features, such as language model embeddings, structural templates, genetic search, and wet-lab experimental data such as contacts determined by cross link mass spectrometry or epitope mapping.

#### 2.1.1 Language model embeddings

Protein structure prediction models like Chai-1 are typically trained with multiple sequence alignments (MSAs) to capture co-evolutionary information. Over the past few years, a number of language models have been introduced and shown to enable accurate prediction of protein structure. However, these models have yet to demonstrate actionable performance on multimeric prediction or protein-ligand interactions. To enable strong single-sequence capabilities in Chai-1, we add an additional input track consisting of residue-level embeddings from a large protein language model [13, 14, 8]. At inference time, we find that this enables accurate prediction across the full suite of tasks.

#### 2.1.2 Constraint features

We also add new training features, designed to mimic experimental constraints. These include pocket, contact, and docking constraints, which capture varying granularity of interactions between entities in a complex.

These work similarly to how templates provide intra-chain distances, but are focused instead on providing inter-chain distance information (see Methods for additional details). These features are trained with dropout to prevent the model from overly relying on these constraints. During inference, these constraints can be specified using prior knowledge or information gained from experiments such as hydrogen-deuterium exchange mass spectrometry or cross-linking mass spectrometry.

### 2.2 Protein-ligand prediction

We evaluate Chai-1 on the PoseBusters benchmark set [15], which measures protein-ligand interactions. All structures in this benchmark set are released on or after 2021-01-13 – after our training data cutoff. We follow the recommendations provided in Buttenschoen et al. [15] and Abramson et al. [11] to compute success rate on this benchmark as the fraction of predictions with ligand root mean square deviation (ligand RMSD) to the ground truth lower than 2 Å. We rank samples by the interface pTM-score of the ligand and penalize ligands that do not respect the input chirality by dividing their interface pTM-score by 100. We do not predict structure 7D6O, which has been marked as obsolete in PDB. For 4/428 structures that have more than 2048 tokens, we crop the assembly to the 2048 tokens closest to the ligand binding site.

Given only the sequence of the protein and the chemical composition of the ligand, Chai-1 achieves a ligand RMSD success rate of 77%, which is comparable to the 76% achieved by AlphaFold3^3^ (Figure 2, Table 1). To evaluate the prompting and conditioning capabilities of Chai-1, we also evaluate on a docking task, which is the original setting proposed by the authors [15]. Namely, specifying the apo structure of the protein boosts success rate to 81%. We note that the holo structure of the protein could leak conformational information that makes the task easier, and we therefore primarily consider this task as a way to evaluate the prompt following abilities of the model.

**Table 1.** Fraction of predictions with ligand RMSD ≤ 2 Å on Posebusters V1 dataset (restricted to 427 structures after removal of PDB ID 7D6O)

| Method | Success rate |
| --- | --- |
| Chai-1 | 77.05 % |
| AF3 | 76.34 % |
| RF2AA | 42 % |
| Chai-1 - Docking | 81.20 % |

**Figure 2.**
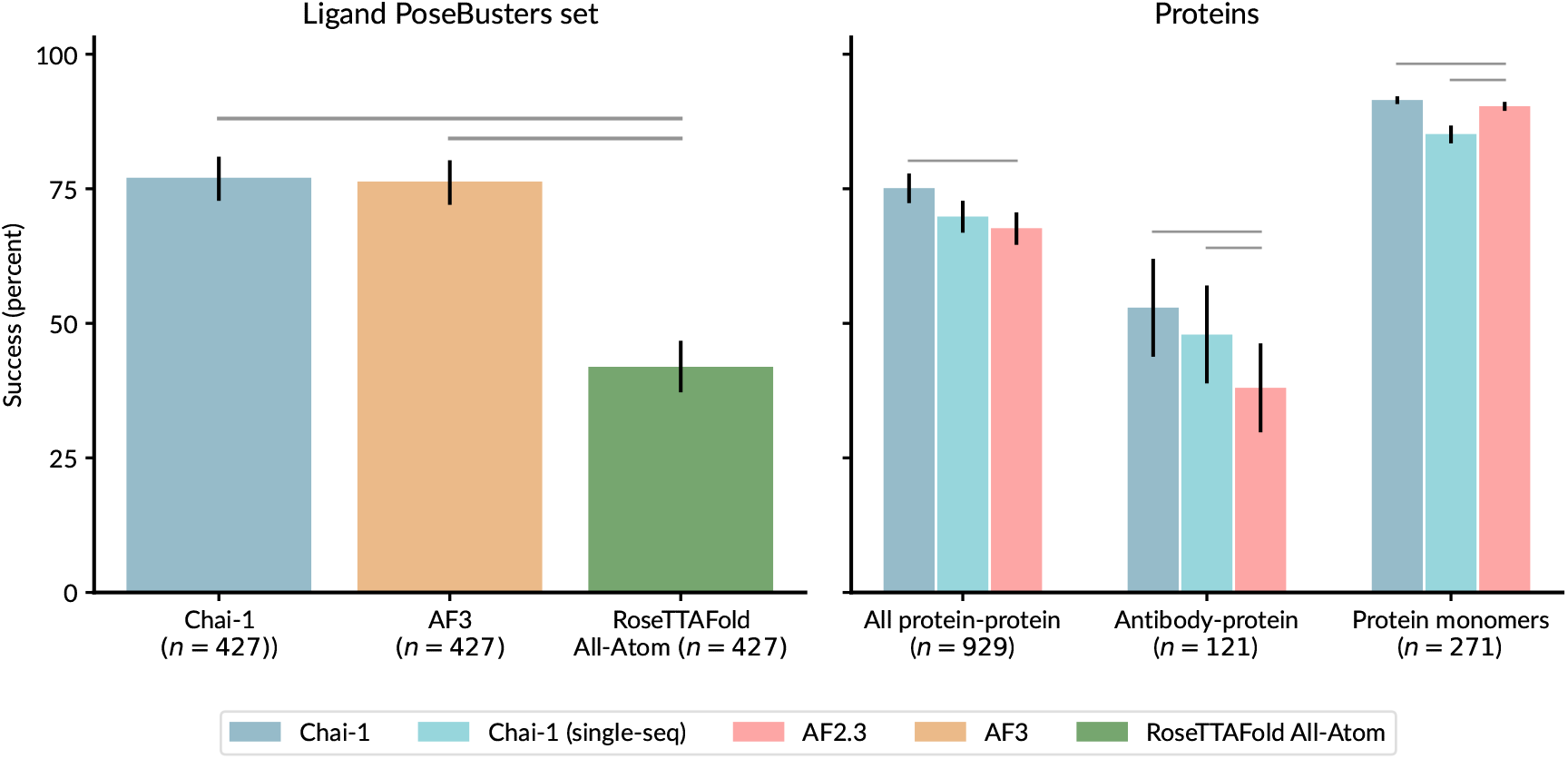
Chai-1 performs at the state-of-the-art across a variety of tasks relevant to drug discovery. Panels show percentage success rates across a variety of modeling tasks. Metrics are: percentage of pocket-aligned RMSD *<* 2 Å for ligands, DockQ *>* 0.23 for protein-protein and antibody-protein interfaces, and C^*α*^-LDDT for protein monomers. Horizontal bars indicate statistically significant differences. Error bars are calculated with an exact binomial distribution for PoseBusters, and 10,000 bootstrap samples for all others.

To aid in future community benchmarking efforts, we make Chai-1 predicted structures of the Posebusters set available. We also provide the results of detailed chemistry checks performed by the Posebusters python package [15] (Supplementary Figure S1).

To explore the limitations of our model and our approach to benchmarking it, we manually inspect examples where our model underperforms (Figure 3). In the case of 7Q2B [16], Chai-1 predicts the ligand as deeper inside the binding pocket than the ground truth structure suggests. However, the ground truth structure also indicates a dimethyl sulfide (DMS) molecule situated deep in the binding pocket. DMS is typically used as a crystallization aid for elucidating structures and its presence may prevent the ligand from binding in a deeper conformation. In the absence of DMS, we find that aromatic ligands from the same campaign [16] appear to be situated deeper in the binding pocket, suggesting that the alternative pose predicted by Chai-1 may indeed be feasible (Figure 3). This highlights the importance of manually inspecting models, and is one of our motivations for releasing the Chai-1 code and model weights.

**Figure 3.**
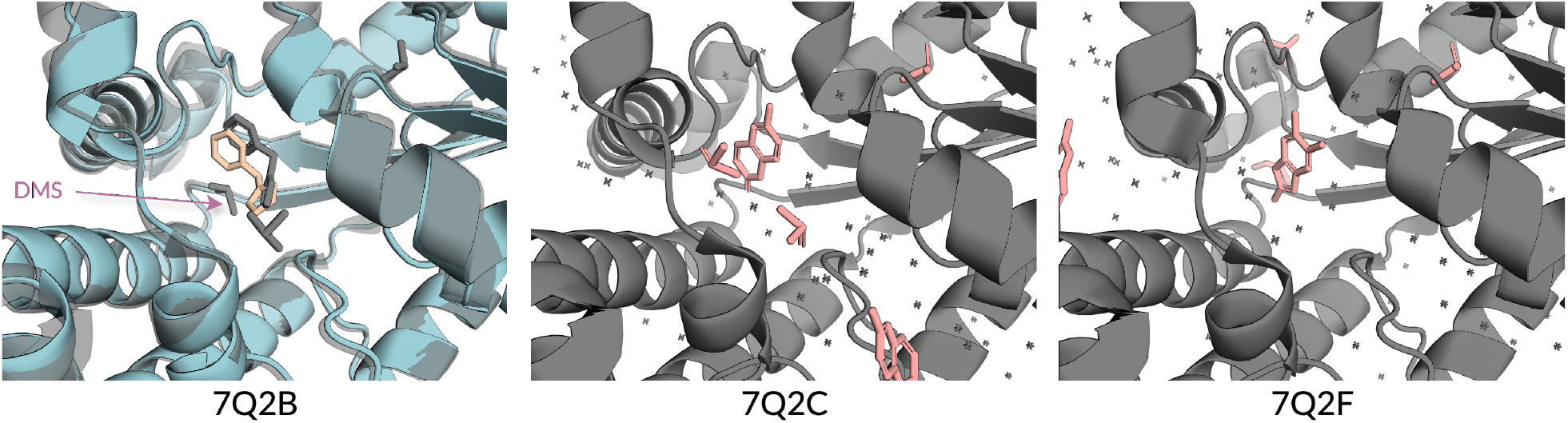
Exploring model and benchmarking limitations. Left panel shows Chai-1’s top-ranked prediction (ligand RMSD 1.73 Å) on the PoseBusters structure 7Q2B, compared to reference ground truth structure (gray). Chai-1 predicts the ligand (tan) deeper inside the pocket (blue) than the ground truth structure; however, the ground truth structure contains a DMS molecule deep in the pocket that may be blocking the ligand. Structures for two similarly aromatic ligands studied in the same work [16] (7Q2C, center; 7Q2F, right) show ligands with two aromatic rings (peach) similar to that present in 7Q2B binding deeper in the pocket (gray) in the absence of DMS.

### 2.3 Multimeric protein prediction

We evaluate performance on protein multimers on an evaluation set comprised of protein structures from the PDB that were released after Chai-1’s training date cutoff. We further filtered this evaluation set to a set of protein-protein interfaces with low homology to our training set (see Methods). This resulted in a set of *n* = 2362 protein-protein interfaces among *n* = 1054 PDB structures, which we then clustered to obtain *n* = 929 interface clusters. We predict the full complex of each structure using Chai-1 and Alphafold Multimer 2.3 (AF2.3) and score the low-homology interfaces using DockQ. To avoid bias in the data distribution, we cluster complexes at 40% sequence identity and report average DockQ success rates (i.e. fraction of predictions with DockQ > 0.23) across clusters (Figure 2). We find that Chai-1 significantly outperforms AF2.3 on this evaluation set (two-sided Wilcoxon test, *p* = 6.24 × 10^−10^, Table 2) with an average success rate of 0.751 compared to 0.677. Without templates, Chai-1 achieves a slightly lower DockQ of 0.743; this reduction is not statistically significant. In single-sequence mode without MSAs, Chai-1 also performs comparably to AF2.3 with MSAs (0.698 vs. 0.677; difference between means is not statistically meaningful). Notably, while previous work has studied single-sequence models for multimer folding [17], prior methods have fallen short of AF2.3 in terms of performance. Chai-1 is the first model to offer high-accuracy multimer folding without the need for MSAs, while also outperforming AF2.3 when using MSAs.

**Table 2.** Protein structure prediction performance on evaluation sets. Leading values indicate mean, values in parentheses indicate 95% confidence interval from 10,000 bootstrap samples. Protein-protein and protein-antibody interfaces are scored by DockQ success rate; protein monomers are scored by C^*α*^-LDDT.

| Method | Protein-protein | Protein-antibody | Protein monomers |
| --- | --- | --- | --- |
| Chai-1 (with MSA) | 0.751 (0.723, 0.778) | 0.529 (0.438, 0.620) | 0.915 (0.907, 0.922) |
| Chai-1 (single-sequence) | 0.698 (0.668, 0.728) | 0.479 (0.388, 0.570) | 0.852 (0.834, 0.867) |
| AlphaFold 2.3 multimer | 0.677 (0.646, 0.706) | 0.380 (0.298, 0.463) | 0.903 (0.895, 0.911) |

Antibodies are an increasingly popular class of therapeutic molecules due to their attractive drug-like properties [18]. We created a subset of the evaluation set that includes only interfaces where one entity was determined to be an antibody (see Methods for details). This set includes 268 interfaces across 129 structures, forming 122 redundancy reduced clusters. On this antibody-protein evaluation set, Chai-1 outperforms AF2.3 by a significant margin, as measured via DockQ success (DockQ > 0.23, two-sided Wilcoxon test, *p* = 3.25 × 10^−5^, Table 2). In fact, Chai-1 without MSAs in single-sequence mode performs similarly to Chai-1 when given full MSAs on these antibody-protein interfaces, and also outperforms AF2.3 (which is still provided MSA information, two-sided Wilcoxon test, *p* = 4.04 × 10^−4^). We hypothesize that MSAs have little performance impact for Chai-1 on this antibody set because there is less evolutionary signal available in the first place for these more variable sequences. This echoes the findings of prior works developing single-sequence folding methods [19]. Following this logic, Chai-1 in single-sequence mode appears to be a particularly potent method for exploring design space of highly variable immunological protein sequences as it is both fast and accurate.

We also repeated the above comparisons with more stringent DockQ cutoffs (Figure S2). We find that Chai-1 maintains its performance lead over AF2.3 even under more stringent thresholds for DockQ success, indicating that Chai-1 improves multimer interface prediction across the board. More broadly, these above results suggest that Chai-1 sets a new benchmark for multimer protein folding in both its full performance mode with MSAs, and its single-sequence mode without MSAs and without structural templates.

We also evaluate Chai-1 on a subset of antibody-antigen protein interfaces. Note that this subset differs from the more general antibody-protein subset as it restricts specifically to antibody-antigen interactions. Furthermore, in cases where multiple copies of an antibody/antigen pair appear in the same pdb, we evaluate independently on each copy. For this reason, the number of interfaces reported here will be larger than that of the general evaluation set. We illustrate the impact of our pocket and contact features by simulating experimentally derived constraints in the form of epitope residues and pairwise distance restraints. In Figure 4 we see that predictions generally improve as more restraint information is provided. When the model is conditioned on a single antibody-antigen distance restraint (*θ* ≤ 15 Å), the percentage of DockQ acceptable predictions increases from the baseline of 35% to 57%. Conditioning on four randomly sampled epitope residues more than doubles the DockQ success rate across all quality cutoffs compared to baseline (Blind) performance. Unfortunately the fraction of high quality predictions still remains low (4-8%) which suggests that high quality antibody-antigen structure prediction remains a generally challenging task.

**Figure 4.**
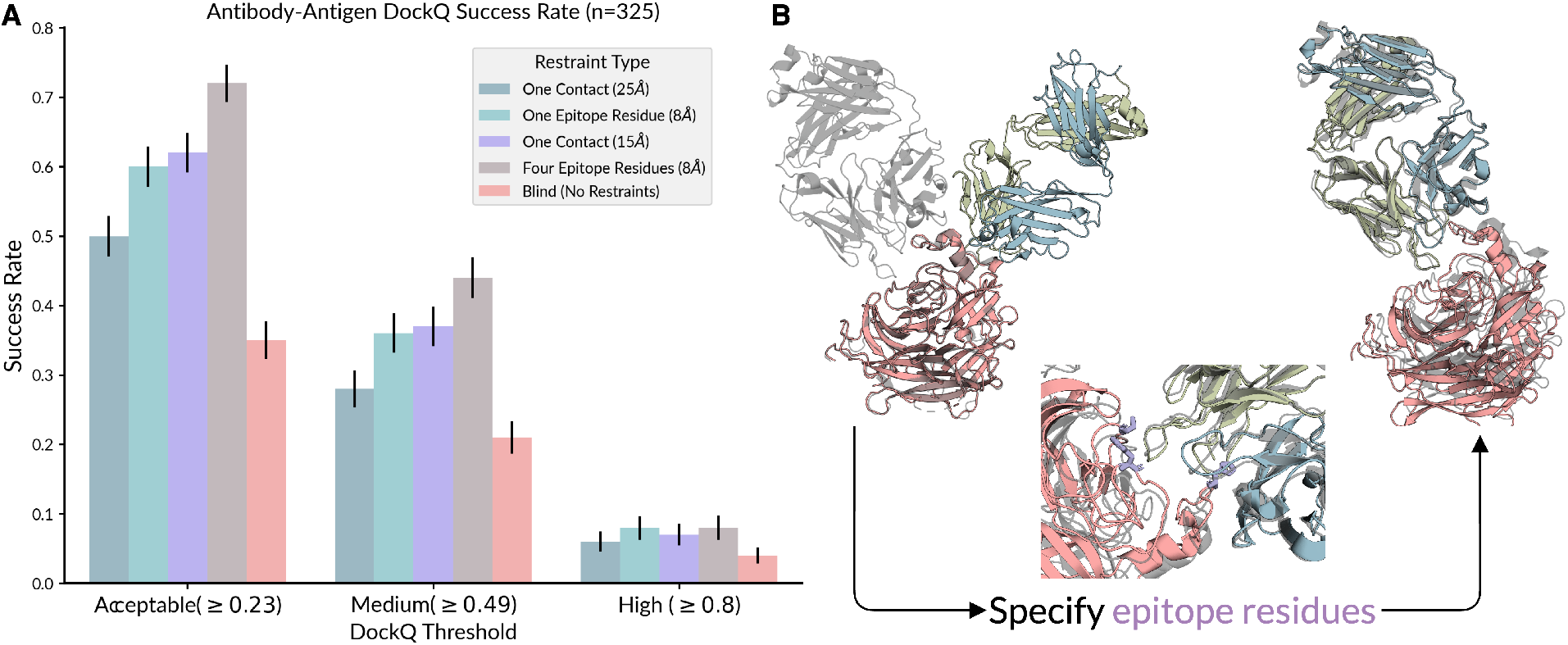
Effect of incorporating docking constraint features on antibody-antigen complex prediction. **A)** DockQ success rates for antibody-antigen interfaces from the low homology evaluation set. DockQ is measured on interfaces where one chain is specified as an antibody and the other as an antigen according to SAbDAb annotation [20]. Bars are grouped by acceptable, medium, and high quality predictions. Bars show performance of Chai-1 in five settings, *One Contact (15A)*/*One Contact (25A)* we randomly sample a pair of residues, one in the antibody chain and one in the antigen chain with distance less than 15Å/25Å respectively and provide the pair and distance threshold to our model as a contact feature. *One Epitope (8 Å)*/*Four Epitope (8 Å)* we sample one/four antigen residue(s) that have minimum C^*α*^ distance less than 8Å to some residue in the antigen chain and provide this residue and chain as a pocket feature. Finally, the *Blind* setting shows our models performance without constraints provided. **B)** Cherry-picked example predictions for PDB ID 7SYV with and without epitope residues (pocket restraints, purple) provided. True structure is shown in gray, predicted chains are colored. Conditioning on epitope residues improves this prediction dramatically increasing the DockQ score from 0.10 to 0.81.

Unfortunately, AF3 restricts commercial use and so we cannot benchmark DeepMind’s latest model on the above protein-protein and protein-antibody evaluations.

### 2.4 Protein monomer prediction

We next curated a low-homology protein monomer set consisting of 447 protein monomers, belonging to 271 clusters. We ran Chai-1 with full MSA information, Chai-1 in single sequence mode, and AF2.3 on this set of protein monomers and evaluated the accuracy of the predicted structures by C^*α*^-LDDT (Figure 2). We find that Chai-1 when given full MSA information outperforms AF2.3 by a small but statistically significant margin (two-sided Wilcoxon test, *p* = 7.21 × 10^−10^). Without MSA information, Chai-1 exhibits poorer performance on monomer folding than AF2.3 (two-sided Wilcoxon test, *p* = 4.46 × 10^−16^).

Due to commercial use restrictions, we are unable to directly compare to several recent methods – including AF3 and ESM3 – for protein structure prediction. We attempt to fairly compare to some of these methods by additionally evaluating Chai-1 on the Critical Assessment of Structure Prediction 15 (CASP15) protein structure prediction targets, consisting of 70 protein monomers. Of these, we exclude one structure that could not complete inference on Chai-1 in a reasonable timeframe (T1169). On the remaining 69 targets, we find that Chai-1 achieves an average LDDT of 0.849, whereas AF2.3 – the previous state-of-the-art monomer folding model – achieves an average LDDT of 0.843. If we focus specifically on the targets that AF2.3 struggles to predict (AF2.3 LDDT < 0.75), we find that Chai-1 predicts significantly more accurate structures for these examples (average LDDT of 0.643 compared to 0.552 for AF2.3, two-sided Wilcoxon test, *n* = 14, *p* = 3.66 × 10^−4^). ESM3 [21] also reports results on the full set of 70 CASP15 targets, and achieves an average LDDT of 0.801 with their largest 98-billion parameter model.

### 2.5 Nucleic acid structure prediction

We evaluate Chai-1 on nucleic acid structure prediction tasks, running the model in single sequence mode (i.e. no RNA MSAs) in all cases. We evaluate performance on a low-homology evaluation set constructed in a similar fashion to our low-homology evaluation sets for proteins and protein-protein interfaces. Owing to the lower number of examples for protein-nucleotide complexes, we report performance metrics averaged across individual examples instead of first averaging within clusters and reporting average performance across clusters. We find that Chai-1 has similar performance to RosettaFold2NA on these complexes as evaluated using interface LDDT (Figure S3). We also compare Chai-1’s performance against that of RoseTTAFold2NA on 9 CASP15 RNA targets, measuring LDDT of C1^*′*^ atoms; we find that the two methods produce comparable results here as well. All nucleic acid results are achieved in spite of the fact that Chai-1 is trained and performs inference without MSAs for nucleic acid sequences, whereas RoseTTAFold2NA has full access to such evolutionary information. Future work incorporating nucleic acid MSAs or nucleic acid language model embeddings [22, 23] could improve its accuracy when modeling these complexes.

As previously stated, we cannot directly evaluate the performance of AF3 on these structures due to commercial use restrictions. AF3 reports a LDDT of 0.473 on 8 of the CASP15 RNA targets, which suggests improved performance on RNA structures. Like RoseTTAFold2NA, AF3 was trained with nucleic acid MSAs, whereas Chai-1 was not.

### 2.6 Predicted confidence scores track accuracy

We find that model confidence estimates from Chai-1 are well calibrated with prediction quality. In Figure 5 we show that interface predicted TM score (ipTM) is a strong discriminator of model quality across all molecular interaction types. We note that the set of interfaces evaluated in Figure 5 include all model predictions rather than only the top prediction for each interface, ranked by confidence. We also note that that proteinligand interactions are restricted to non-bonded ligands and exclude ions.

**Figure 5.**
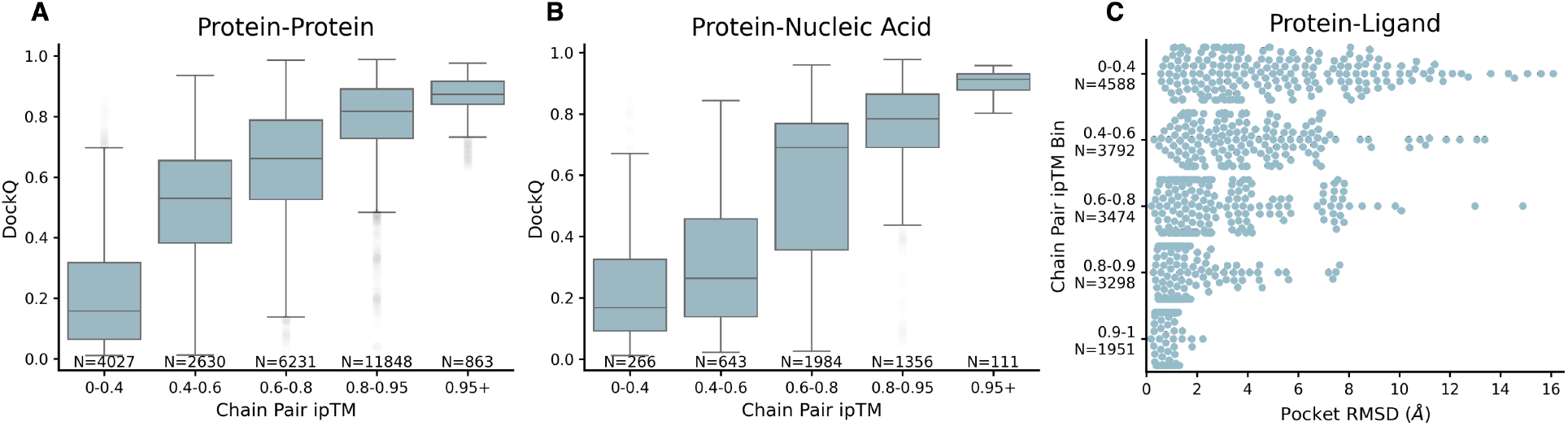
Assessment of model predicted confidence scores against ground truth structures on interfaces from the evaluation set. Here we show results for all interfaces in our low homology subset, and do not restrict only to interfaces with low homology. All pdbs in this set have a later PDB release date than our training set cutoff. Chain-pair ipTM is shown for each plot and refers to the ipTM score of the two chains defining the interface being evaluated. **A)** Boxplot of DockQ against binned chain pair ipTM restricted to protein-protein interfaces. **B)** Boxplot of DockQ against binned chain pair ipTM restricted to protein-nucleic acid interfaces. **C)** Swarm plot of pocket RMSD against binned chain pair ipTM. Dots show ligand RMSD, sampled down to 5% of the overall data. Note that Protein-Ligand interactions are restricted to non-bonded ligands and exclude ions.

### 2.7 Chai-1 lab server

The Chai-1 lab server uses the Chai-1 Github repository as a dependency. To enable low-latency MSA generation, we developed a distributed MSA search pipeline which runs jackhmmer in parallel on multiple shards of the genetic databases UniRef90, Uniprot and Mgnify. Unlike our training and evaluation data, the server does not query the Uniclust30+BFD database or use the `-N 3` flag on jackhmmer. We spot-checked the difference in results between the server and our internal inference pipeline on a set of examples from the Posebusters evaluation set, finding that they differ by *<* 1Å RMSD in all cases (*µ* = 0.345Å, σ = 0.150Å), even though the accelerated server version was run without templates. We note that these changes may still result in subtle discrepancies between the server results and those in the paper, and we therefore refer readers to the GitHub repository where all variables can be controlled.

We provide a series of example structures predicted by the Chai-1 lab server in Figure 6. These have been chosen to emphasize potentially therapeutically relevant structures. We show the Zika virus NS2B-NS3 protease in complex with a small molecule inhibitor ligand (7VLH, Huber et al. [24]), a SARS-CoV-2 protease in complex with a non-structural protein fragment (7MB4, Shaqra et al. [25]), and a complex containing PD-1 and the monoclonal antibody checkpoint inhibitor cemiplimab (7WVM, Lu et al. [26]).

**Figure 6.**
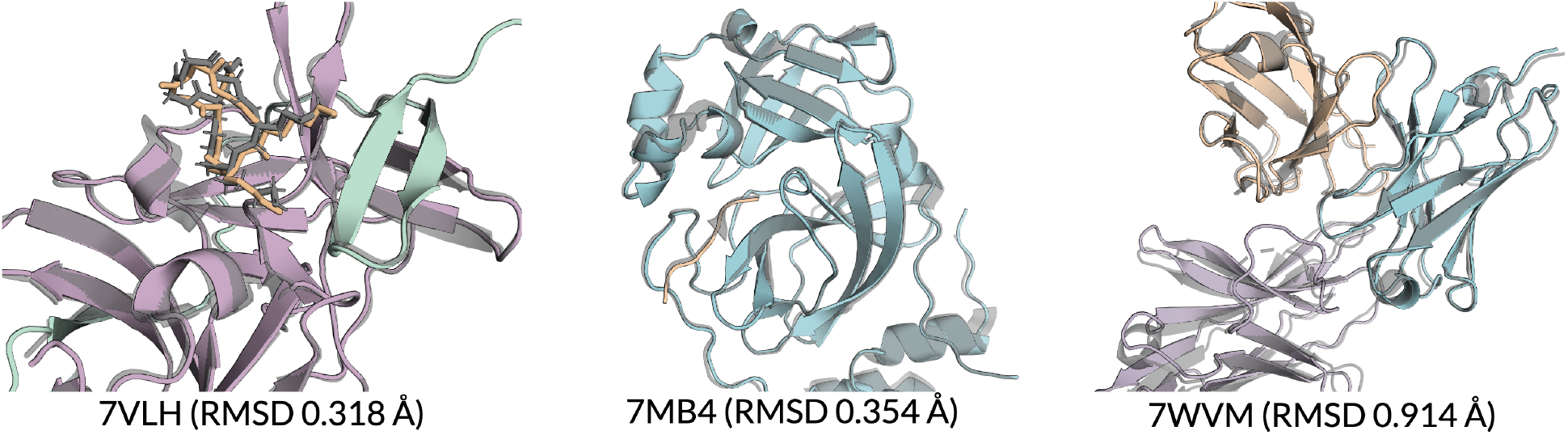
Examples of predictions from Chai-1 lab server. Each panel shows Chai-1 lab server predictions on a complex released after Chai-1’s training cutoff date. MSAs are included for all examples, though the procedure for generating them differs from our main results. Ground truth structures are shown in gray. Predicted chains are shown with various colors; one color corresponds to each chain in the complex. RMSD values comparing the ground truth and Chai-1’s outputs are shown under each panel.

### 2.8 Limitations

A limitation we frequently encountering when evaluating Chai-1’s predictions is that the model may sometimes predict the individual chains in a complex correctly, but fail to place them in the correct relative orientations. An example of this behavior is shown in the first panel of Figure 4, where the predicted complex is poor without additional contact information.

Another notable limitation is that Chai-1 can be highly sensitive to modified residues. Removing modified residues from a sequence that natively posses them or replacing modified amino acids with their standard amino acid analogs can cause large changes in predicted structures. We hypothesize that this is because Chai-1 has been trained explicitly on structures with modifications and relies on this information to accurately predict structures. More plainly, those same amino acid sequences without modifications might be considered to be entirely different inputs.

## 3 Discussion

We believe that building an accurate understanding of the structure of biological molecules is foundational to advancing our scientific understanding of cellular processes, and ultimately, for advancing human health. In this spirit, we are excited to share our latest folding model, Chai-1. We are excited to build and improve this model with the greater scientific community.

## 4 Contributors

Jacques Boitreaud, Jack Dent, Matthew McPartlon, Joshua Meier, Vinicius Reis, Alex Rogozhnikov, Kevin Wu.

## 5 Methods

### 5.1 Architecture

The Chai-1 neural architecture largely follows previous work Abramson et al. [11], with a heavy reliance on pair-bias self-attention. Key differences to previous work is summarized below.

#### 5.1.1 Language model embeddings

Chai-1 is trained on a combination of multiple sequence alignments and protein language model embeddings generated by a 3 billion parameter model [8]. This model generates per-residue embeddings for each input protein sequence. Modified residues are replaced with their canonical “parent” residue if available, and are otherwise replaced with an unknown (“X”) amino acid type. Residues that do not belong to a protein chain (i.e. DNA, RNA, or ligands) are assigned a mask token.

At inference time, Chai-1 can be run with any combination of MSAs, templates, and language model embeddings.

#### 5.1.2 Constraints features

Pocket constraints are represented by a token ID, *i*, chain ID, *C*, and distance threshold, *θ*_*P*_. The feature specifies that min_*j*∈*C*_ ∥*x*_*i*_ − *x*_*j*_∥ ≤ *θ*_*P*_. During training, we randomly sample *θ*_*P*_ ∈ (6, 20) and randomly sample satisfying pocket constraints from the ground truth structure. Contact constraints are represented by a pair of tokens *i* and *j*, and a distance threshold *θ*. Similar to the pocket constraint, we randomly sample *θ*_*D*_ ∈ (6Å, 30Å) and satisfying constraints ∥*x*_*i*_ − *x*_*j*_∥ ≤ *θ*_*D*_ from the ground truth structure. The docking feature is a one-hot encoding of pairwise distances between subsets of input tokens using four bins [0−4Å, 4−8Å, 8−16Å, *>* 16Å]. This differs from template information, as while templates contain intra-chain distance information, they do not contain inter-chain distances. Docking constraint features are generated by first partitioning the chains of the ground truth complex into two groups, and then generating pairwise distances between all token center coordinates within each group.

During training, we also apply chain-wise and token-wise dropout on constraints features, i.e. we sometimes provide docking constraints for only a subset of the tokens and chains in the true structure. Additionally, each of these features is included independently with probability 10% during training. When a feature is not included, a separate learnable mask value is used. In order to facilitate accurate prediction in a variety of constraint settings, we also randomly sample the number of distance and pocket restraints according to a geometric distribution with parameter 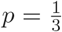.

### 5.2 Data and Training

We trained Chai-1 on 128 Nvidia A100 GPUs with a batch size of 128 for 30 days. Chai-1 is trained on both PDB and AlphaFoldDB [27] structures. For PDB structures, we apply with a release date cutoff of 2021-01-12. We do not train on any of the distillation datasets described in [11] other than AFDB. Templates are generated using PDB70 with the same cutoff date. We largely follow the data preprocessing and clustering steps from Abramson et al. [11].

#### 5.2.1 Genetic search

Whenever available, we use MSAs that have been generated and deposited in OpenProteinSet [28] for databases UniRef90 [29], UniProt [30], MGnify [31], and UniClust30+BFD [6]. Following [11], when such MSAs are not available, we compute MSAs using `jackhmmer v3.4` [32] on the UniRef90, UniProt, and MG-nify databases, with the arguments `-N 1 -E 0.0001 --incE 0.0001 --F1 0.0005 --F2 0.00005 --F3 0.0000005` as well as an additional `--seq_limit` flag that is set to the following for each database:

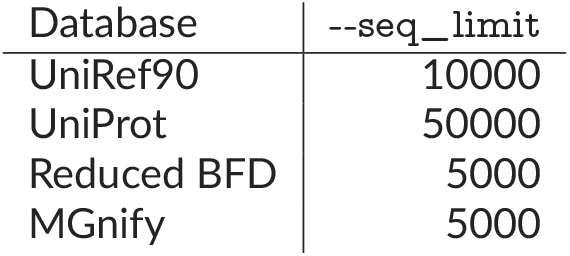

| Database | --seq_limit |
| --- | --- |
| UniRef90 | 10000 |
| UniProt | 50000 |
| Reduced BFD | 5000 |
| MGNify | 5000 |

For inference over the datasets in our main analysis (CASP15, the low-homology evaluation sets, and Pose-Busters), we additionally generated MSAs using `jackhmmer` with 3 iterations (`-N 3` flag) on the UniRef90, UniProt, and MGnify databases. All databases use the same date cutoffs and versions described in [6]. To avoid excessive computational time, the web interface only provides MSA with `n=1` JackHMMer iteration, although the open-source code can be run with any user-provided MSA.

### 5.3 Inference

Unlike Abramson et al. [11], we use the same model for all evaluations since our training data cutoff does not overlap with the data used in any of our evaluation sets. We run Chai-1 with 4 recycles, 5 trunk samples and 5 diffusion samples for a total of 25 predicted structures unless otherwise noted. A confidence model is then used for ranking (details below). We run inference by giving Chai-1 the input sequence, MSAs, protein language model embeddings, and templates. In single sequence mode, we provide only the input sequence and protein language model embeddings; MSAs and templates are omitted in this mode.

Unless otherwise noted, inference is run on FASTA and/or SMILES sequences derived from the first biological assembly in the corresponding target .cif file. In all cases we remove crystallization aids, mirroring how our model was trained. Inference is currently limited to a maximum of 2048 tokens.

### 5.4 Baseline methods

We ran AlphaFold-multimer 2.3 [12] on protein monomers and multimers via the popular community wrapper ColabFold [33], version 1.5.5. ColabFold uses the same model weights as AlphaFold, but notably replaces the expensive genetic search procedure for building MSAs with a faster search via MMseqs2 [34]. ColabFold has been shown to be significantly faster than the original AlphaFold2 code without performance regressions [33], and outperforms the original AlphaFold pipeline on the CASP15 blind structure prediction assessment (https://predictioncenter.org/casp15/zscores_final.cgi). We follow the ColabFold MSA generation documentation, and as input to ColabFold, we provide MSAs generated by running MMSeqs2 against UniRef30 (version 2303) and `colabfold_envdb` (version 202108). We do not search for or provide templates, as prior works have shown minimal effect on model performance (indeed, two of the five AlphaFold2 models were not trained on templates in the first place). We use weights for AlphaFold2-multimer version 2.3, running with 3 recycles across 5 models and 5 seeds, and without relaxation. We use the top-ranked output across these 25 examples. We verified that this setup produces results largely concordant with values present in the literature. Namely, Hayes et al. [21] reports that AlphaFold2 achieves 0.826 LDDT on *n* = 70 structures in CASP15. In our reproduction, we obtain an LDDT of 0.842.This value differs slightly from the value we discuss in our results section, as it includes all 70 structures whereas the main results exclude one structure.

We ran RoseTTAFold2NA via Tamarind Bio’s API https://www.tamarind.bio/api-docs, which also follows the best practices and open-source codebase for that method. All other performance results are taken from their respective original works. We are unable to evaluate several methods, including AF3 and ESM3, due to commercial use restrictions.

### 5.5 Metrics

Protein-ligand complex prediction is evaluated using pocket-aligned ligand RMSD. This is computed by first aligning the predicted and native structures using interface atoms in the native structure. Interface atoms are defined as any C^*α*^ atom within 10 Å of any ligand atom in the true structure. After alignment, RMSD is computed over ligand atoms to calculate pocket-aligned ligand RMSD.

We measure the quality of protein-protein interfaces using DockQ [35]. When comparing complexes that require fewer than 8! = 40320 chain mapping permutations to exhaustively enumerate, we enumerate all permutations and select the chain mapping permutation that optimizes global DockQ score averaged across all interfaces. For larger complexes, we adopt an alignment-based strategy as follows. For each predicted chain ĉ, we consider all native structure chains *c* that share the same sequence. We choose a native chain *c*_*j*_ find the transformation that, when applied to the predicted chain ĉ_*i*_ minimizes RMSD between ĉ_*i*_ and *c*_*j*_.

We then apply this transformation to the entire predicted complex *C* and greedily assign a chain mapping by selecting the nearest predict chain for each native chain. We generate such chain mappings considering all possible combinations of ĉ_*i*_ and *c*_*j*_ and select the chain mapping permutation that optimizes global DockQ score averaged across all interfaces. In both cases, we evaluate individual interface DockQ scores within the overall complex. Note that this approach differs significantly from the simulated annealing approach described by Abramson et al. [11]. Our approach avoids inconsistent chain assignments that may arise from randomness and local minima associated with the simulated annealing process.

We use the local difference delta test [36] (LDDT) computed over reference atoms to evaluate folding accuracy on protein and nucleotide monomers. Protein LDDT is evaluated with an inclusion radius of 15 Å on C^*α*^ atoms. An inclusion radius of 30 Å nucleotides is used on C1^*′*^ atoms. Protein-nucleotide interfaces are evaluated using interface LDDT (iLDDT) over C^*α*^ and C1^*′*^ atoms in the interface.

### 5.6 Evaluation set

Our evaluation set is constructed from a temporal split of PDB entries released between 2022-05-01 and 2023-01-12; these structures were all released after any data in our training set (most recent cutoff date of 2021-01-12). We additionally restricted to non-NMR structures with resolution better than 4.5 Å. We restricted to examples that had 2048 or fewer tokens. From this dataset, we removed monomers and interfaces with substantial homology to the training set:

- Monomers with 40% or greater sequence identity to the training set are removed.
- Polymer-polymer interfaces where both polymers have greater than 40% sequence identity to two chains in the same complex in the training set are filtered out.
- Polymer-peptide interfaces where the non-peptide entity has 40% or greater sequence identity to the training set are removed.

We additionally cluster the above low-homology evaluation set. Individual polymer chains were clustered at 40% sequence identity for proteins more than 9 residues, and 100% sequence identity for proteins with 9 or fewer residues and for all nucleic acids. Clustering on interfaces is accomplished by assigning a cluster ID to each participant in the interface, and constructing a full interface cluster id from those components:

- Polymer-polymer interfaces are given a cluster ID of (polymer1_cluster, polymer2_cluster).
- Polymer-ligand interfaces are assigned the cluster ID of the polymer entity.

We defined the antibody-protein set as all interfaces where one of the two chains in the interface belong to one of the two largest clusters in the evaluation set. We found these to be chain B in the PDB structure 7PKL, and chain C in 7T2P – corresponding to light and heavy chains, respectively.

Evaluation on this dataset is done on either individual monomer chains, or on specific low-homology interfaces extracted from a full complex prediction. Note that in the case of complexes, the overall complex that we predict may contain chains that have significant homology to the training set, but we only evaluate on interfaces that are low-homology as defined above. We use Biological Assembly 1 for all evaluation.

We note that while the above methodology for constructing an evaluation set follows the procedure laid out in Abramson et al. [11], we are not able to match exactly the evaluation set used in AF3. As a result, values reported on protein-protein interactions, protein-antibody interactions, protein-DNA and protein-RNA interactions are not directly comparable to those reported in Abramson et al. [11].

## 6 Supplementary Information

### 6.1 Supplementary tables and figures

**Figure S1.**
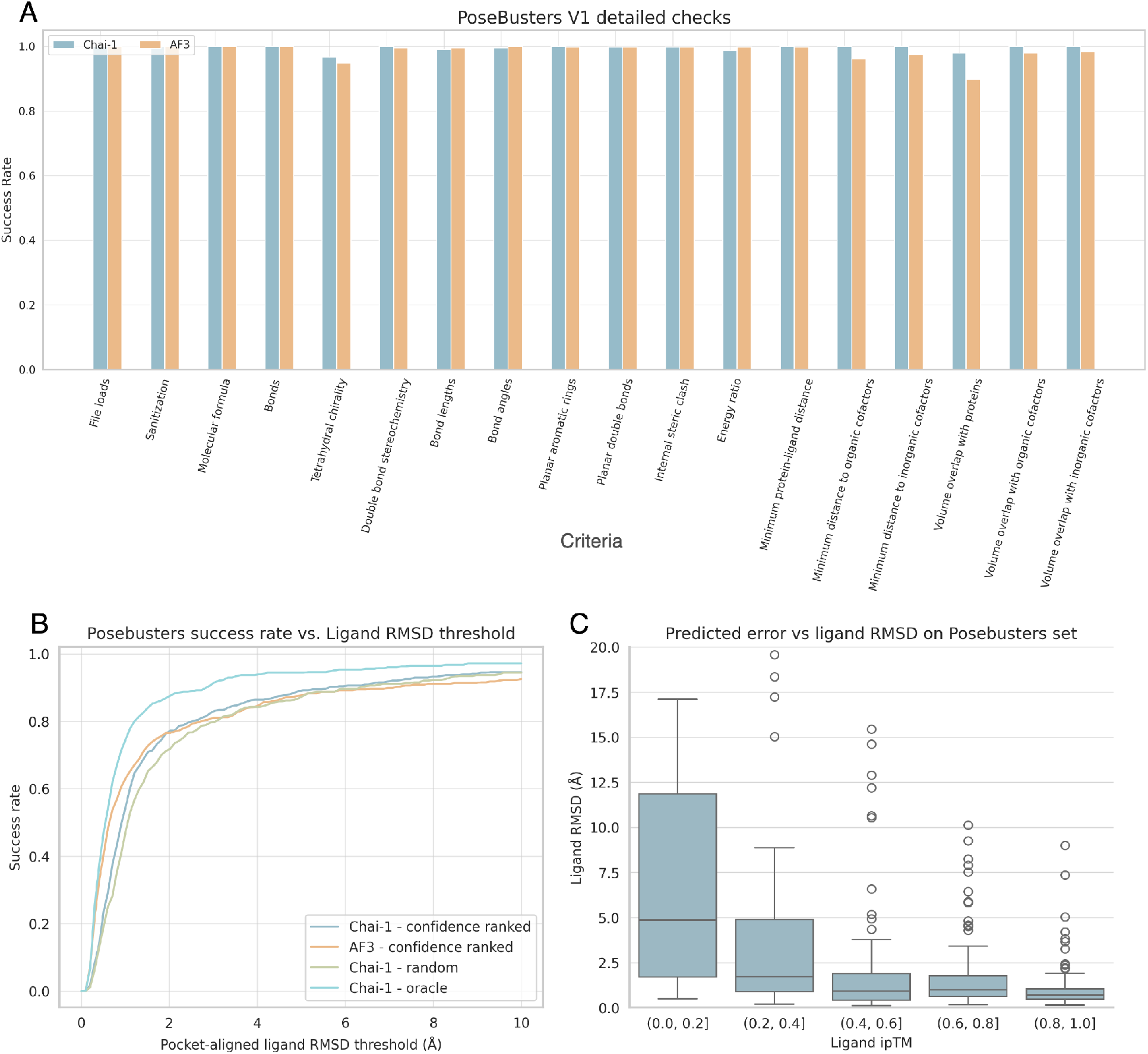
Extended data for posebusters evaluation. **A)** Posebusters detailed checks on predicted structures for Chai-1 and AF3. Chai-1 generates chemically valid structures, but sometimes struggle to respect the chirality of the input ligand, an issue also observed in AF3. Notably, Chai-1’s predicted structures exhibit a lower inter-molecular clash rate than AF3; we did not have to use a clash penalty for sample ranking. **B)** Cumulative density plot of ligand RMSD of model predictions on Posebusters set. Chai-1 confidence ranked takes the best sample among 5 trunk samples and 5 diffusion samples, ranked by ligand ipTM. Chai-1 random takes the mean over the aforementioend 25 samples, and Chai-1 oracle takes the best (lowest ligand RMSD) prediction after comparing all 25 examples to the ground truth. **C)** Binned ligand ipTM confidence score against top-ranked prediction ligand RMSD on the Posebusters evaluation set.

**Figure S2.**
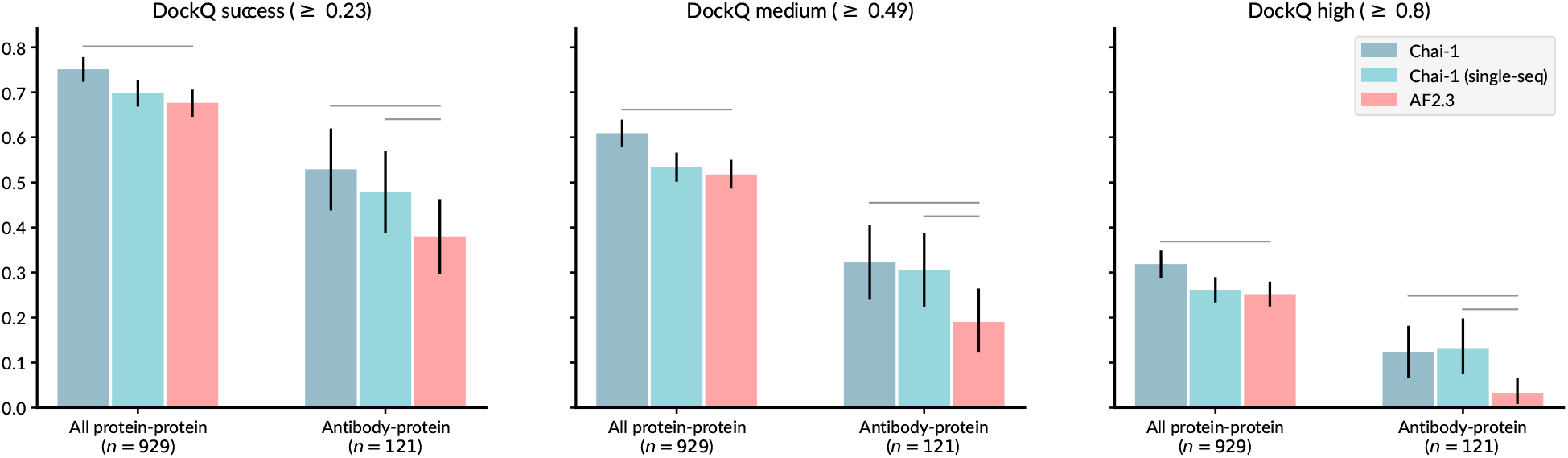
Chai-1 protein-protein DockQ success rates with increasing cutoffs. Each panel shows the DockQ success rate for acceptable, medium, and high quality predictions (left to right, cutoffs for each shown in title). Error bars are computed with 10,000 bootstrap samples. Horizontal bars indicate statistically significant differences with a two-sided Wilcoxon test.

**Figure S3.**
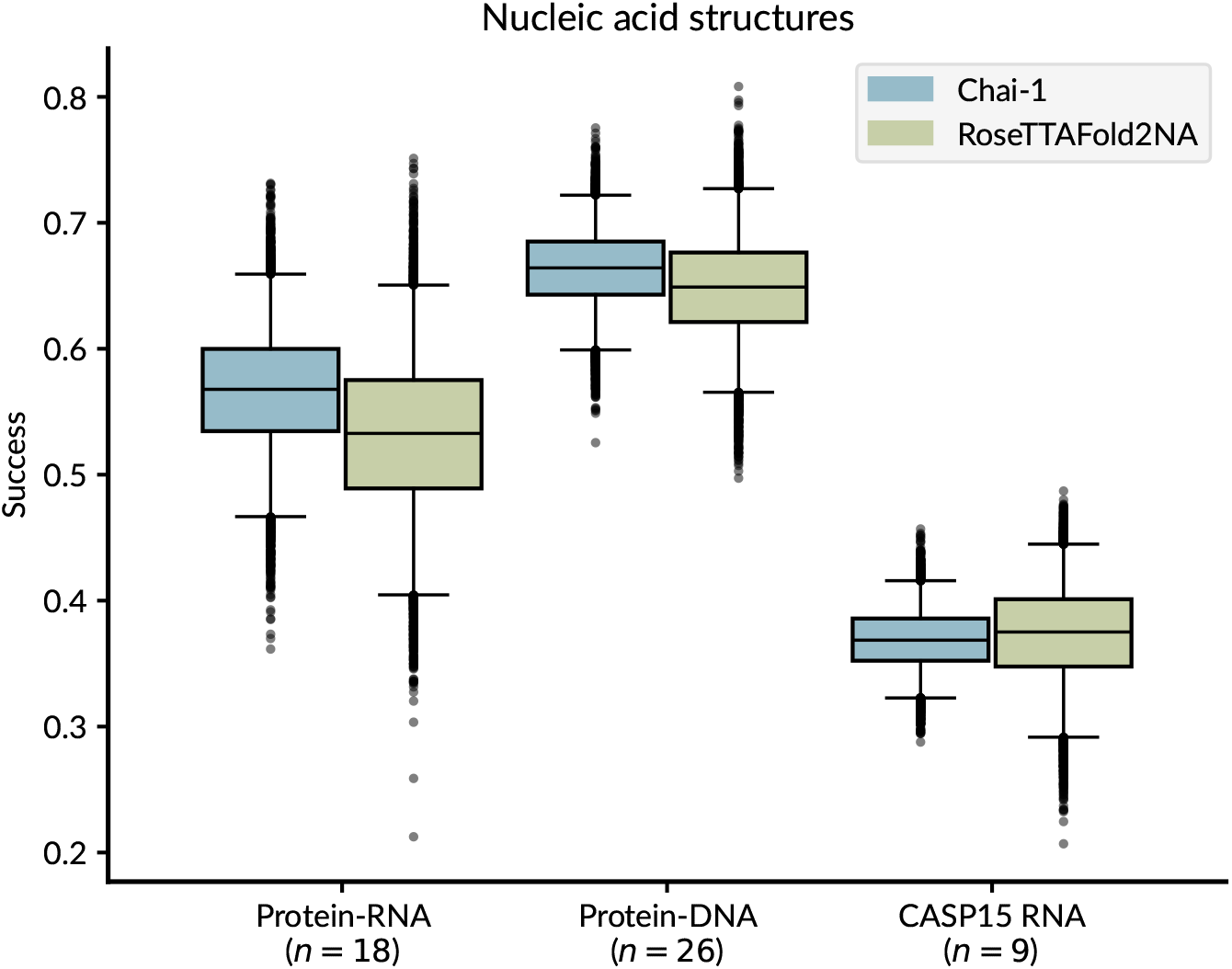
Chai-1 performance on nucleic acid complex prediction. Protein-nucleic acids complexes are evaluated using interface LDDT, and RNA complexes are evaluated using C1^*′*^-LDDT. Boxes indicate the inter-quartile range across 10,000 bootstrap samples, and whiskers indicate 95% confidence interval (also across 10,000 bootstrap samples).

### 6.2 Supplementary data

Predicted structures for the posebusters set are available at https://chaiassets.com/chai-1/paper/assets/posebusters_predictions.zip

## Footnotes

1 https://github.com/chaidiscovery/chai-lab/

2 https://lab.chaidiscovery.com/

3 Values are taken from AlphaFold3’s publicly released PoseBusters predictions; we did not run AF3 ourselves.

